# A Computational Pipeline to Identify and Characterize Binding Sites and Interacting Chemotypes in SARS-CoV-2

**DOI:** 10.1101/2022.03.24.485222

**Authors:** Sarah H. Sandholtz, Jeffrey A. Drocco, Adam T. Zemla, Marisa W. Torres, Mary S. Silva, Jonathan E. Allen

## Abstract

Minimizing the human and economic costs of the COVID-19 pandemic and of future pandemics requires the ability to develop and deploy effective treatments for novel pathogens as soon as possible after they emerge. To this end, we introduce a unique, computational pipeline for the rapid identification and characterization of binding sites in the proteins of novel viruses as well as the core chemical components with which these sites interact. We combine molecular-level structural modeling of proteins with clustering and cheminformatic techniques in a computationally efficient manner. Similarities between our results, experimental data, and other computational studies provide support for the effectiveness of our predictive framework. While we present here a demonstration of our tool on SARS-CoV-2, our process is generalizable and can be applied to any new virus, as long as either experimentally solved structures for its proteins are available or sufficiently accurate homology models can be constructed.

## 2 Introduction

To minimize the human and economic costs of a pandemic, effective treatments for novel pathogens like SARS-CoV-2 must be developed as quickly as possible. While drug target selection has improved over time, it remains difficult, and poor target selection often leads to early failures in the drug development process. An accurate and efficient computational method for identifying and characterizing protein binding pockets (**i.e.** potential drug target sites), in emerging pathogens would thus serve a critical purpose in shortening the timeframe for drug development by providing more viable starting points for experimental screenings. To enable a timely and rapid response to a new pathogen, such target identification techniques must be adaptable to unfamiliar disease agents.

Molecular docking simulations, Molecular Mechanics Generalized Born Surface Area (MM/GBSA) methods, and molecular dynamics simulations have traditionally been employed to gauge the nature and quality of proteinligand binding and have been performed broadly in the study of SARS-CoV-2 proteins.^1–7^ The webserver D3Targets-2019-nCoV, for example, uses molecular docking to predict potential drug targets and drug compounds specifically for COVID-19.^8^ Though such simulations provide detailed insight of physicochemical properties at the atomistic level, they are computationally costly and limited to short timescales. A variety of computational platforms have been designed to minimize or circumvent these drawbacks.

Existing platforms for binding pocket detection in proteins rely on a range of strategies, broadly based on templates, geometry, physicochemical properties, or machine learning. The PDBspheres program^9^ draws on experimental measurements of protein-ligand complex structures from Protein Data Bank (PDB) to detect and assess potential binding pockets through superposition of structural models. The resulting PDBspheres data includes the set of protein residues predicted to be in contact with each ligand as well as a set of descriptive metrics. Other hybrid approaches combine methods from the four broad categories above. For instance, dPredGB^10^ couples a geometric detection method with the calculation of desolvation properties, while PocketMatch^11^ and SiteHopper^12^ integrate a representation of the surface shape with a description of chemical properties. The “binding response descriptor” is based on a sphere method, refined with clustering, and energy and geometry calculations from docking simulations.^13^ Similarly, Pockets 2.0^7^ features the direct integration of docking simulations into Fpocket. Methods like these that incorporate physicochemical properties through calculations or simulations include an additional perspective in their description of protein binding pockets, but they also depend on predictive *in silico* data, which have not necessarily been validated experimentally and whose quality and accuracy may vary across different pockets. Still other approaches, including WaveGeoMap,^14^ Deeply - Tough,^15^ and Site2Vec^16^ make use of neural networks. While such deep learning techniques enable scalability to enormously large data sets, they create a layer of abstraction between input data and results, making them less interpretable. These platforms lay a foundation for protein binding pocket detection tasks but also leave room for new hybrid approaches that offer both template/geometry-based and machine learning-based methods – roots in purely structural experimental data with a capability to ingest large quantities of input data in a straight-forward manner.

Building on this body of work, we introduce a unique computational framework for the rapid identification and characterization of binding pockets in the proteins of SARS-CoV-2 and other novel pathogens. Our new quantitative and automated approach takes into account all available experimental structural data from PDB to uncover the combination of significant residues and core ligand components. Our goal is to distinguish and characterize what we call “consensus pockets,” a term we put forward both as a general concept and a more narrow, practical definition for the specific task in this work. Conceptually, consensus pockets could refer to structurally similar pockets from different organisms (structural consensus across organisms) or to pockets within a single organism that bind similar classes of ligands (consensus of ligand features), for example. The overarching notion of consensus pockets is relevant to a variety of applications and when tailored appropriately, defines the key factors of interest in a given application. For instance, structural consensus across different viruses would be important for developing multi-target drugs, while ligand consensus or structural consensus between viral and human proteins would be critical for identifying and avoiding off-target effects of prospective drug candidates. For this particular application, we define consensus pockets as those binding pockets with the strongest consensus of contacts across different protein-ligand complexes, based on the residues in contact with experimentally observed or computationally predicted binding ligands and certain features of those ligands (described in more detail in the next section).

To this end, we first combine molecular-level structural modeling of protein-ligand binding and pocket detection from PDBspheres^9^ with a hierarchical clustering procedure to determine and describe relevant binding pockets. While protein-ligand binding data from any pocket detection platform could serve as input to our framework, we choose to use data from PDBspheres because the metrics it reports enable more comprehensive and refined pocket characterization, the primary goal of this work. We then employ cheminformatics tools and density-based clustering to uncover the pocket-specific, key chemical substructures involved in binding. Though our framework relies on access to protein structural data which may vary in quality, we account for such differences by incorporating into our analysis a measure of the quality of our structural models. We also apply several strategies for substantiating our results and find strong qualitative and quantitative support.

Capable of producing results in a matter of days, our framework is designed to act as a rapid response tool in the face of an emerging pathogen. Our novel approach enables cross-organism analysis of pocket structural similarities and can reveal which, if any, binding pockets in a new pathogen are more conserved, which could aid drug discovery efforts for broad-spectrum antivirals.^17,18^ The combination of residue and ligand clustering provides a more well-rounded perspective and is a notable feature of our process. In establishing a link between protein pockets (and even pocket subregions) and key chemical features, it allows us to make focused suggestions of molecular structures that are better suited for specific binding pockets. Additionally, the relative simplicity of our system makes it more straightforward and interpretable. Taking advantage of the fact that structural and physicochemical information is inherent in PDB entries and hence in PDBspheres data, our pipeline uses minimal parameters. Most importantly, our framework is generalizable. Provided there are experimentally solved structures or homology models of sufficient quality, it can be applied to any new virus, or more broadly, to any class of proteins with structurally similar binding pockets.

## 3 Methods

The foundation of our approach lies in molecular-level knowledge of protein-ligand binding, which we obtain through experimental data from PDB and protein structural modeling from PDBspheres. ^9^ Experimental measurements of protein-ligand structures from Protein Data Bank (PDB) serve as the input to PDBspheres, which uses superposition of structural models to detect and evaluate potential binding pockets. The Global Distance Calculation (GDC) metric^19,20^ takes both the backbone and side chains into account and is a measure of the structural similarity between pairs of PDBspheres structural models. For a given protein-ligand binding complex, the higher the calculated GDC score, the greater the structural similarity between the PDBspheres models of the SARS-CoV-2 protein of interest and the reference ligand binding pocket observed in experimentally solved structures deposited in PDB. The approach presented here relies on the PDBspheres data set of predicted residue-ligand contacts for SARS-CoV-2 proteins, which was generated using the entire PDBspheres library of approximately two million templates to ensure that as many binding sites as possible were detected. Using the GDC metric as a means to evaluate our data and understand our results, we leverage the residue-level resolution of PDBspheres data to identify relevant binding sites in SARS-CoV-2 and the corresponding site-specific chemical components, as shown in the schematic of our workflow in Fig. 1.

**Figure 1:**
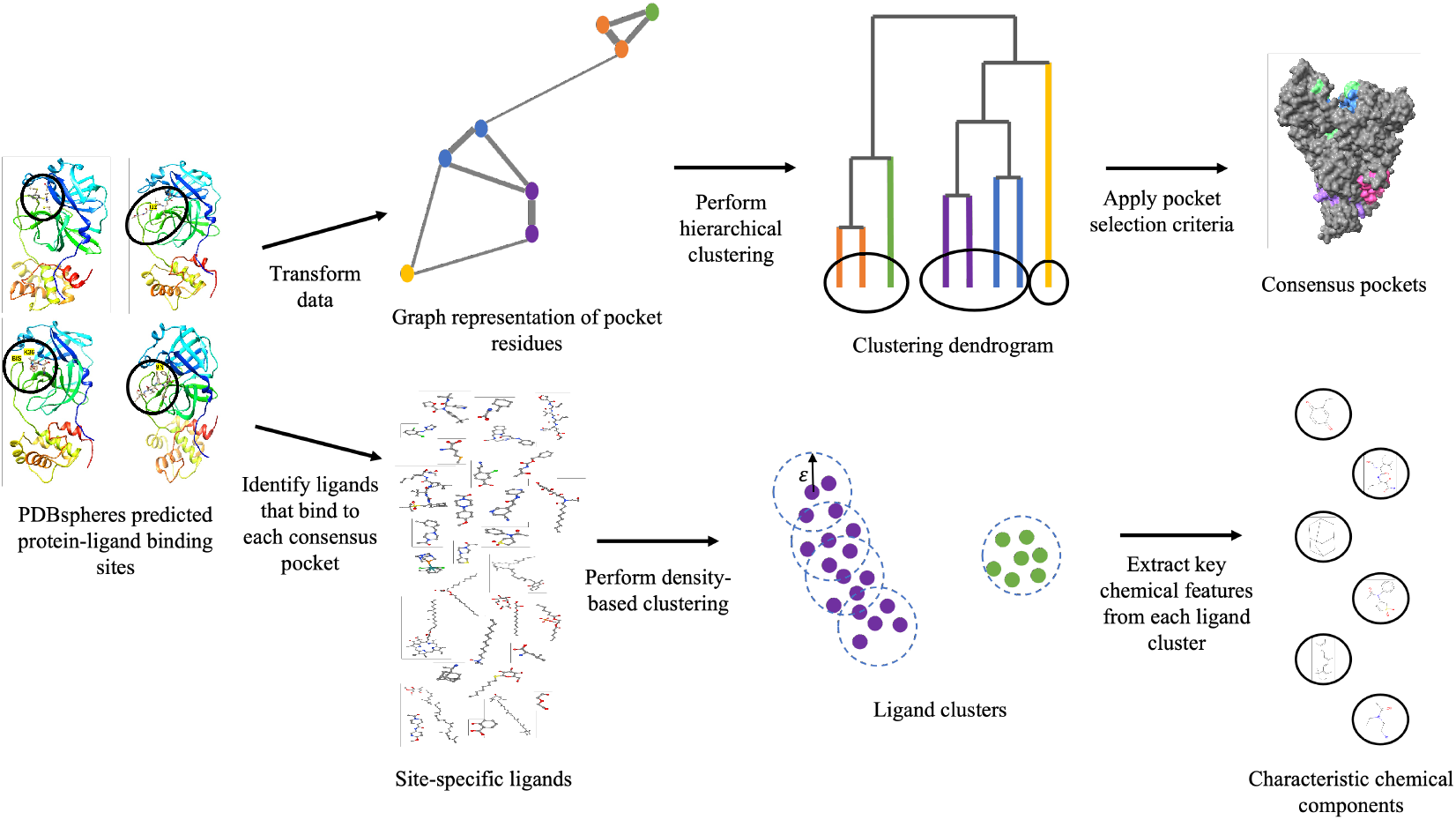
Schematic of our process for identifying relevant binding pockets (top row) and site-specific chemical components (bottom row).

We begin by assessing the confidence level of all protein-ligand binding pockets detected by PDBspheres, focusing on the quality of evaluated protein models and the relevance of ligands. For some target proteins, there exist experimentally solved, high resolution X-ray structures that are representative of the protein. For other target proteins, only lower resolution structures, fragmented structures, or homology models are available. This range in the quality of available structural models impacts our confidence in the detection of protein-ligand binding pockets and the specific residues in contact with ligands. To estimate this confidence, a set of criteria and thresholds has been introduced in the PDBspheres system. ^9^ In our approach, we use only PDBspheres data that meets the thresholds specified below. We also apply four ligand-specific exclusion criteria.

With the remaining PDBspheres residue-ligand contact data, we use NetworkX ^21^ to construct a weighted, undirected graph for each SARS-CoV-2 protein, in which each node corresponds to a protein residue and each edge indicates that the two connected residues bind one or more of the same ligands (Fig. 1).

From the graph, we calculate pairwise distances between residues (as given by Eqn. 3 below). Based on these calculated distances, we then perform complete linkage hierarchical clustering to group protein residues. The result is a full dendrogram, like that in Fig. 1, indicating which residues or residue groups are clustered together and at what distances those clusters occur. Different distance cutoffs in the dendrogram correspond to different numbers and identities of clusters (*i.e*. putative binding pockets). Given that binding pockets may range in size, as ligands do, they may appear at different length scales in the dendrogram, and we implement a selection process (described below) to identify all relevant binding pockets.

The specific steps we take are as follows:

1. Filter the PDBspheres data based on the criteria below:
  - *GDC* ≥ 60
  - Number of conserved residues *Nc* ≥ 15
  - Number of residues forming an interface with the ligand *N*4 ≥ 4
  - Number of residues potentially clashing with the ligand *cl* = 0
  - For practical reasons, exclude data for ligands whose SMILES string is missing or cannot be parsed.
  - To aid in distinguishing between regions of the protein with different binding behavior, exclude data for non-specific ligands that bind over a third of the residues in a given viral protein.
  - Exclude data for ligands that do not contain at least eight non-hydrogen atoms, which are too small to be relevant.
  - To focus on ligands whose composition may be more relevant for drug discovery, exclude ligands that do not contain at least two of the following atoms: carbon, nitrogen, and oxygen.
2. Construct a weighted, undirected graph for each SARS-CoV-2 protein, in which each node corresponds to a protein residue and each edge indicates that the two connected residues bind one or more of the same ligands.
  - The edge *E_rs_*, connecting residues *r* and *s*, will have an edge weight *w* given by

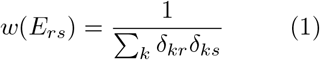

where *δ_kr_* and *δ_ks_* indicate the existence of a contact between ligand *k* and residue *r* or residue *s*, respectively.
  - The summation is performed over all ligands in PDBSpheres.
  - The criterion for a ligand-residue contact is defined as follows:

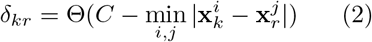

where Θ is the Heaviside step function, 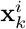 denotes the position of atom *i* in ligand *k*, and 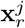 denotes the position of atom *j* in residue *r*.
  - For this calculation, water atoms are excluded from consideration, and the contact threshold *C* = 4.5*Å*.
3. Calculate the distance between pairs of residues.
  1. The distance between two residues is given by the minimum sum of edge weights over all possible paths connecting them. That is, if *P* is a path containing a set of edges connecting residues *r* and *s*, then the distance between residues *r* and *s* is given by

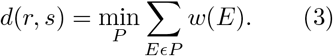
4. Perform complete linkage hierarchical clustering based on the distances given by Eqn. 3.
5. Make sequential cuts over the resulting dendrogram from bottom to top in distance increments of 0.001. Along the way, select as consensus pockets the largest unique clusters that meet the following two criteria:
  - Contain a mean number of contacts per residue of at least 90. This threshold is based on the observed level of noise in the distribution of contacts per residue (Fig. S1)
  - Consist of a biophysically reasonable number of residues - at least 10 and no more than the maximum number of residues in contact with any ligand for the given viral protein

After selecting consensus pockets in each SARS-CoV-2 protein, we perform another clustering routine to identify the site-specific, core chemical components involved in binding. For the ligands in the PDBspheres data, we extract SMILES strings from PDBE Chem and chemical taxonomy information (chemical Kingdom, Superclass, Class, and Subclass) from the Classy-Fire database.^22^ Using techniques for natural language processing, we then create for each ligand a word vector embedding to further identify similarities between ligands. We retrieved the PubMed Central Open Access Subset of biomedical literature^23^ and joined frequent n-grams using the word2phrase tool in four iterations.^24^ We then used the Gensim implementation of Word2vec to train a word embedding with 3,600 dimensions, a 6-token window of optimization, and a minimum token frequency of 5 instances within the corpus.^25^ We compute a vector representing each PDBe ligand using the mean embedding of all synonyms for the chemical compound as listed in the PubChem database.^26^ An embedding was successfully obtained for 1,156 of the 5,180 small molecule ligands used in our PDBspheres analysis; other ligands did not have sufficient frequency in the corpus to estimate an embedding. From the same filtered PDBspheres data mentioned above, we pick out the ligands that bind to each consensus pocket and calculate the following three distances for each pair of ligands *a* and b: the Tanimoto distance *T_ab_*, the chemical taxonomy distance *C_ab_*, and the word vector embedding distance *V_ab_*. We calculate the Tanimoto distance using RDKit, ^27^ representing each ligand as a Morgan fingerprint with a radius of 2 and a size of 1024 bits. We define 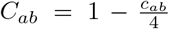, where *c_ab_* is the number of taxonomic levels in common between ligands *a* and *b* and *c_ab_* ∈ [0, 4]. The word vector embedding distance is the cosine distance between vectors *A* and *B* for ligands *a* and b, or 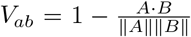. We construct an overall distance metric *W_ab_* as the sum of the available, normalized distances, weighted evenly. For example, if *T_ab_*, *C_ab_*, and *V_ab_* are all available for ligands *a* and *b*, then 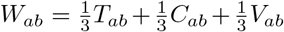. If only *T_ab_* and *C_ab_* are available, then 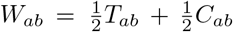. We use a composite distance metric in order to incorporate contextual information about the ligands, allowing us to compare them based on more information than their structures alone. We then cluster the ligands using the Density-Based Spatial Clustering of Applications with Noise (DBSCAN) algorithm ^28^ and our precomputed distance metric. For each binding pocket, we select the parameters *ϵ* and *min_points* by performing a grid search over the values *ϵ* ∈ [1, 1.05, 1.10, 1.15,…, 15] and *min_points* ∈ [1, 2, 3,…, 10] and choosing the combination that yields the optimal clustering, as evaluated by the silhouette score.

## 4 Results and Discussion

Using the PDBspheres data and residue clustering procedure described above, we identify as relevant binding sites the consensus pockets visualized in Fig. 2 for nsp5, nsp12, and Spike. The fact that groups of residues obtained from the residue clustering procedure appear to colocalize in 3D space, as expected, provides confirmation that our method performs as intended. Lists of the constituent residues for each consensus pocket can be found in the table in Fig. 3 for nsp5 and in Tables S2 and S7 in the Supplemental Section for nsp12 and Spike, respectively. The number of consensus pockets and the nature of those pockets can be used to characterize each viral protein. The bar charts in Fig. 3 (for nsp5) and Fig. S3 (for nsp12 and Spike) provide such an initial characterization, showing the number of consensus pockets identified in each protein, the number of residues comprising each consensus pocket (blue), and the number of ligands that bind to each consensus pocket (orange). For example, nsp5 contains one moderately-sized pocket that binds to an extraordinarily high number of ligands (top panel of Fig. 3).

**Figure 2:**
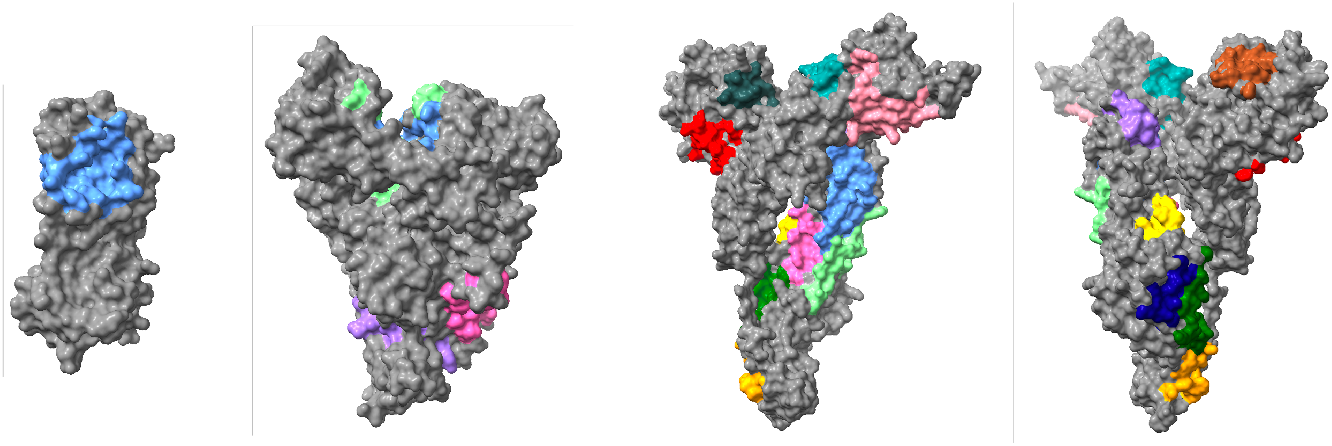
Consensus binding pockets for nsp5 (far left), nsp12 (center left), and Spike (center right and far right). The protein surfaces are colored gray, and within a given protein, each consensus pocket is highlighted in a different color.

**Figure 3:**
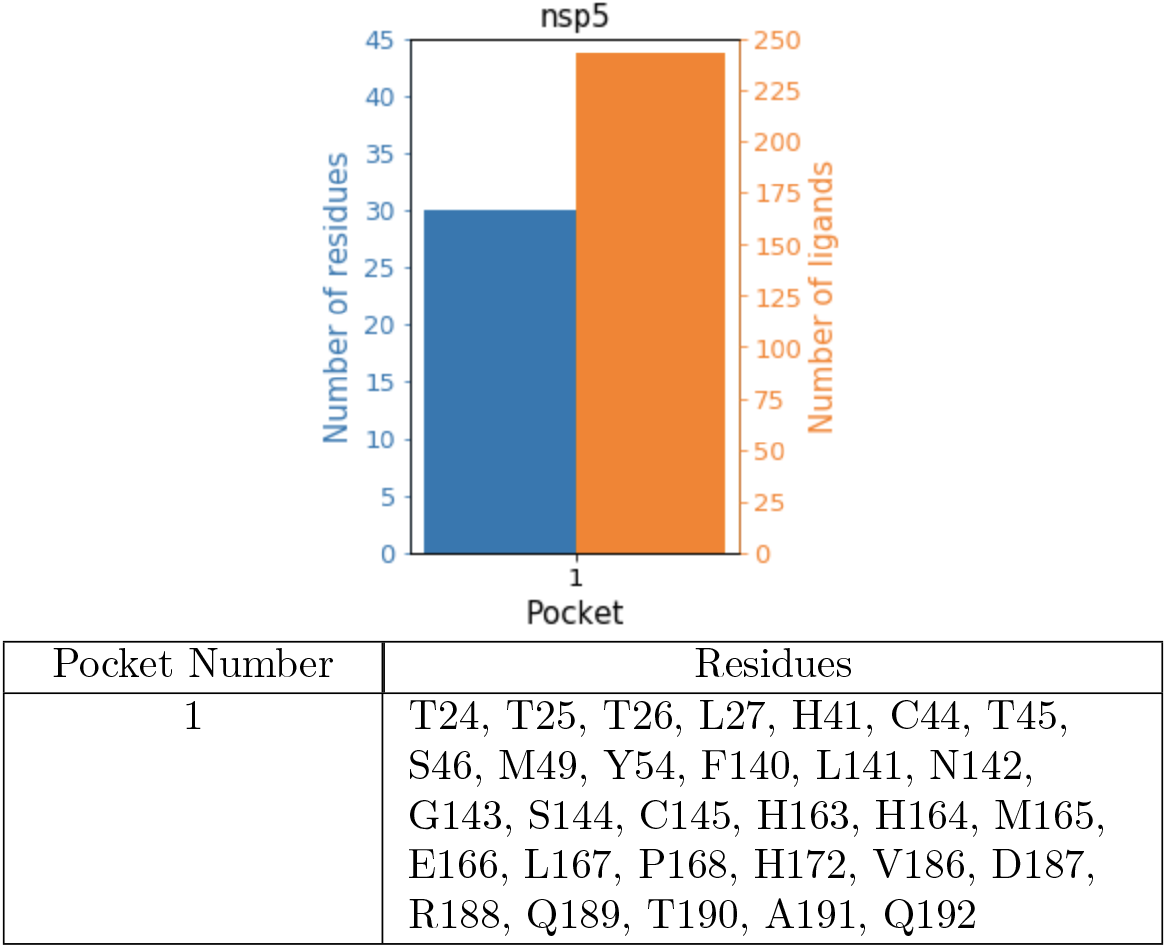
On the top, the bar chart shows the number of constituent residues (blue) and number of ligands (orange) that bind at least 50% of the residues for the consensus pocket in nsp5. On the bottom, consensus pockets for nsp5. All residues belong to the A chain.

Previously published results from both experimental and computational studies corroborate many of the consensus pockets we identify in SARS-CoV-2 proteins. To ensure that the assessment of our method’s performance is fair and draws on different data than was used in its development, we exclude as reference points published pockets derived from PDB entries that are used as SARS-CoV-2 protein models in PDBspheres. For Spike, we identify the same pocket as in Ref. 29. The pocket reported in Ref. 30 and Ref. 2 is only identified by our method when the threshold for the required mean number of contacts per residue is lowered significantly.

As shown in Fig. 4, several of our consensus pockets for Spike, nsp5, nsp12, and nsp16 share a considerable number of residues with previously published pockets. We include zeros in the Venn diagrams in Fig. 4 (and later in Fig. 6) to show lack of discrepancies. Although we do not detect the drug binding pockets highlighted in Ref. 17 for nsp3, nsp13, and nsp16, we do capture most of the same pockets for nsp5, nsp12, nsp14, nsp15, and Spike. Results from PDBspheres indicate that several ligands are predicted to bind to nsp9, and residues in contact with those ligands overlap with 13 pocket residues reported for nsp9;^17^ our workflow described in Methods, however, does not detect any consensus pockets in nsp9 at the currently selected thresholds.

**Figure 4:**
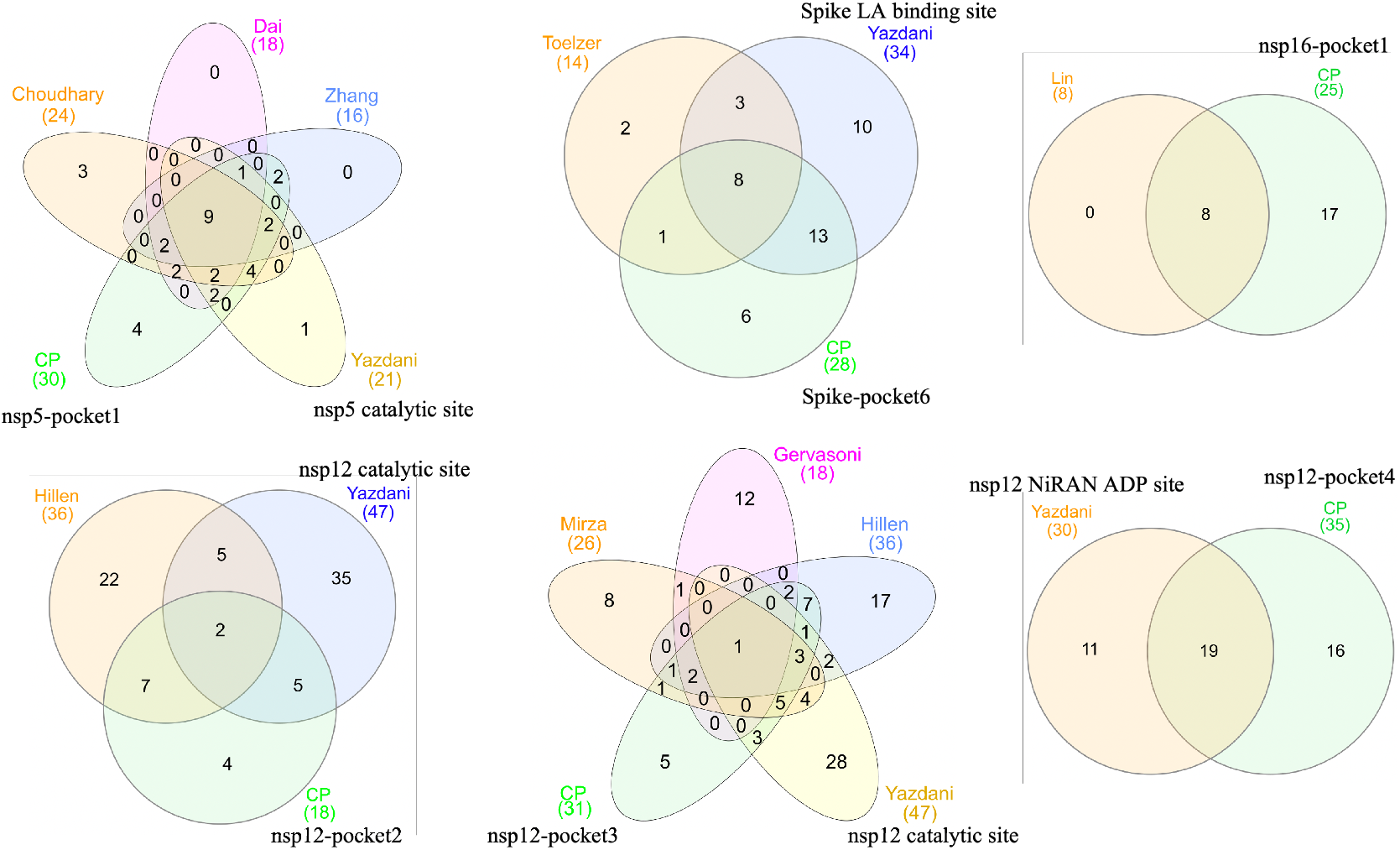
Overlap between our consensus pockets (sets of residues identified by our approach that are predicted by PDBspheres to contact many of the same ligands) and pockets from particular publications (sets of residues identified by the authors as being in contact with a given ligand or ligands). Each colored circle/ellipse represents a different pocket. Numbers indicate the sizes of each pocket and the sizes of the overlaps between various pockets. Circles/ellipses for published pockets are labeled with the last name of the first author, and consensus pockets are colored green and labeled “CP.” Details on published pockets can be found in Ref. 5, 31, 32, 17, 4, 7, 33, 29, and 34.

**Figure 5:**
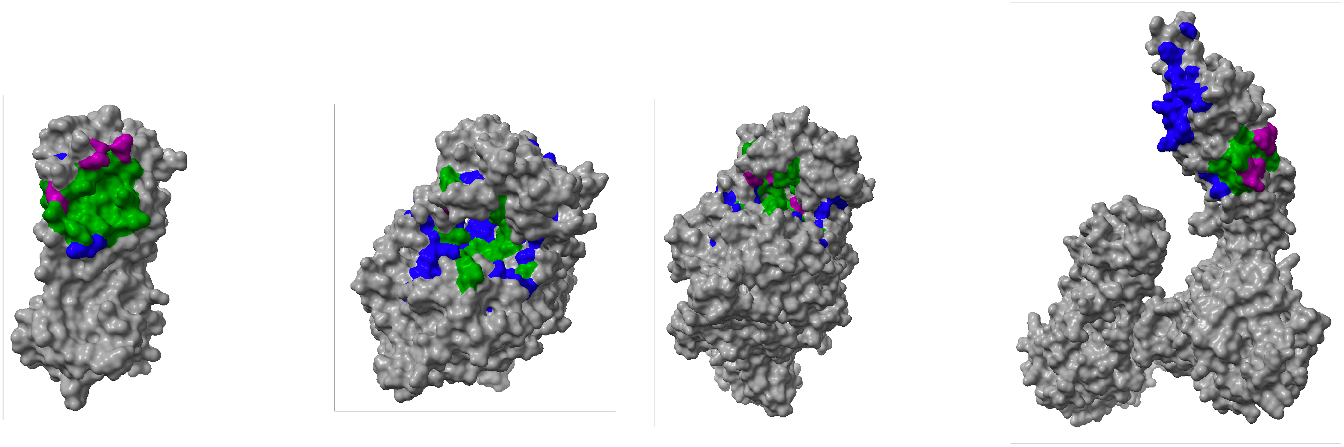
Overlap between published pockets and our consensus pockets for nsp5-pocket1 (far left), nsp12-pocket3 (center left and center right), and Spike-pocket6 (far right). Residues colored blue, purple, and green appear only in published pockets, only in our consensus pocket, or in both, respectively. The results from multiple papers are aggregated for each published pocket. The two images for nsp12 give different views of the same pocket.

**Figure 6:**
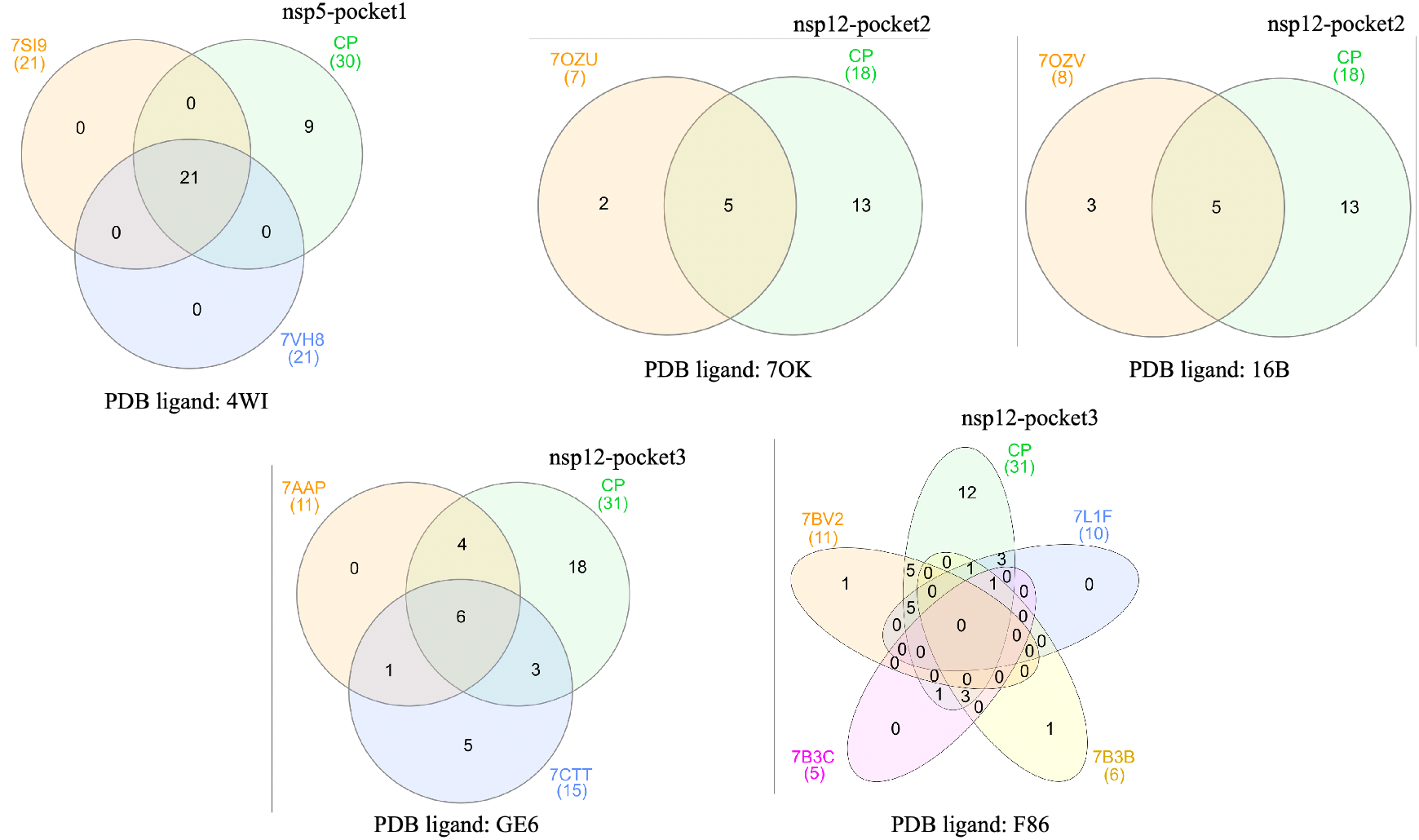
Overlap between pockets from experimentally solved protein-ligand complexes deposited in PDB (sets of residues observed to be in contact with a given SARS-CoV-2 drug or drug form) and our consensus pockets (sets of residues identified by our approach that are predicted by PDBspheres to contact other ligands). PDBspheres data on these SARS-CoV-2 drugs was excluded from the determination of our consensus pockets and saved for a test comparison (displayed in this figure). The PDB ID for the relevant drug or drug form is shown at the bottom of each Venn diagram. The green circle/ellipse in each diagram represents the consensus pocket most closely aligned with the PDB experimental contact residues represented by the other circles/ellipses, which are labeled by their PDB template IDs.

Figure 5 provides a visualization of the comparison in Fig. 4 for some of our consensus pockets. Residues belonging to a given pocket are aggregated across all published papers mentioned in the preceding two paragraphs and are colored according to whether they appear only in the literature (blue), only in our results (purple), or in both (green). The moderately sized green regions in nsp12 and Spike show substantial overlap between our results and those in previously published papers. Notably, the pocket in nsp5 is almost entirely green with only small spots of blue and purple, indicating an especially high degree of overlap between our results and others’ and thus reinforcing the effectiveness of our approach. Overall, while the performance of our consensus binding pocket identification is somewhat mixed for Spike, we clearly and consistently identify the same pockets for nsp5 and nsp12 that are found in other work.

In order to validate our process, we excluded PDB data for known SARS-CoV-2 drugs (those that are currently being tested or have already been approved - at the time of this analysis) before running our pipeline, saving this data as a test set to which to compare our results. Even withholding this data, our consensus pock-ets still include many of the same residues that are in contact with those drugs. Figure 6 shows the number of residues in common between our consensus pockets and experimentally measured contact residues from the test set. A residue is considered to be in contact with a drug if at least one of its atoms comes within 4.5A of any of the drug’s atoms, based on the coordinates in the PDB file. For the held out clinically tested SARS-CoV-2 drugs with PDB structures shown in Fig. 6, each consensus pocket represents a substantial fraction of the contact residues defining the functional pocket. The consensus pockets cover over 50% of the contact residues for all ten templates and often a much higher percentage, up to 100% for four templates. Cases in which there are differences between PDB structural templates, for example 7CTT and 7AAP (lower left diagram in Fig. 6), highlight the natural biological variation in protein conformations that introduces changes to the binding pocket. ^35^ Since the consensus pockets are defined by a comprehensive search of over multiple protein conformations and multiple ligands, the consensus pockets cover the natural variability and still report a majority of residues in contact with antiviral drugs, plus additional residues of potential importance. The fact that our method can discern significant binding pockets without some of the original data is another indication that our method is performing well and could be trusted to identify significant conserved pockets in other new viruses.

Comparison of our consensus pockets against the protein domain families in the Pfam database^36^ provides additional support for our method. Every consensus pocket we identify in nsp5, nsp12, and Spike corresponds to a known Pfam family (see Table 1), suggesting that our process selects relevant sites with functional significance. The results in Table 1 indicate that our consensus pockets are associated with some Pfam families more than others and do not represent a random distribution of Pfam families. Likewise, the distribution of Pfam families for previously published pockets (mentioned above) is also not random, consistent with our results. Based on these observations, it is possible that certain Pfam families are more or less likely to be significant binding sites. The fact that our consensus pockets align with known protein domain families reaffirms that our procedure identifies relevant pockets. In addition, as noted in Pfam, ^36^ the S1 N-terminal domain and the S1 receptor binding domain play a role in binding to host receptors. That many of our consensus pockets for Spike map to these two integral Pfam domains is further evidence that our process selects relevant pockets with an important viral function.

**Table 1:**
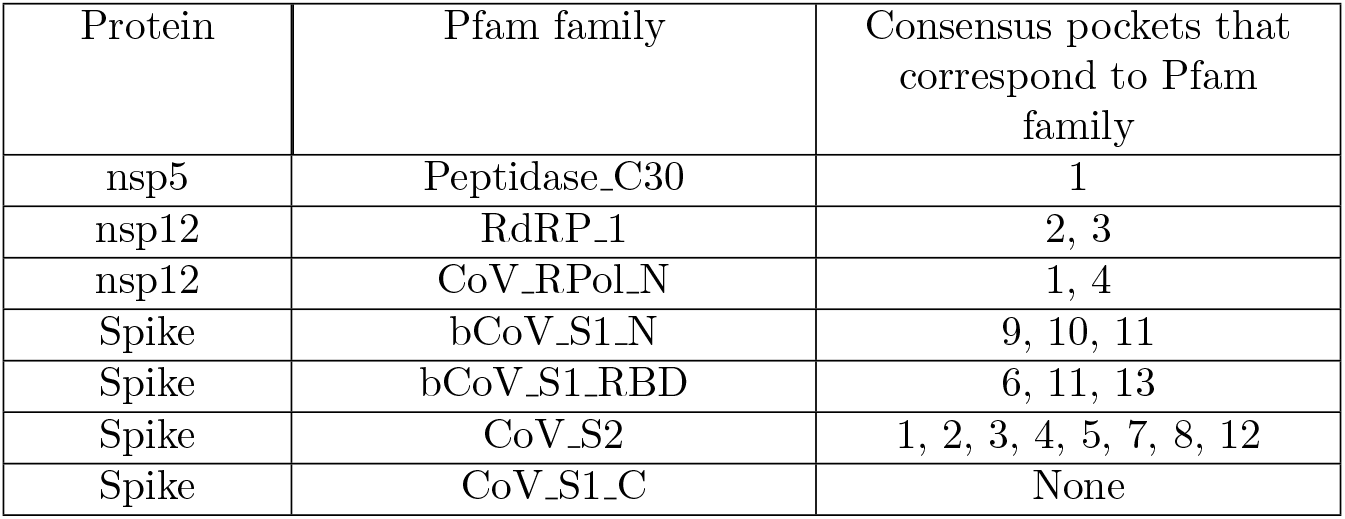
All Pfam^36^ families for nsp5, nsp12, and Spike are shown with the overlapping consensus pockets. All consensus pockets mapped to a Pfam family.

We test the sensitivity of our consensus binding pocket identification approach to the input data by rerunning our residue clustering procedure using different subsets of the PDBspheres data and comparing the resulting consensus pockets. We filter the PDBspheres data by date of deposition of the original data in PDB and the GDC score, which allows us to examine how the passage of time (**i.e.** data availability for a new pathogen), the availability of relevant structural protein-ligand templates, and the accuracy of structural models affect confidence in pocket identification and the overall performance of our approach. The adjusted rand index (ARI) serves as a metric for evaluating the similarity of the consensus pockets across different subsets of the data. An ARI of one indicates that two sets of clusters (**i.e.** consensus pockets) are identical, while an ARI of zero indicates that the clusterings have no clusters with common elements (**i.e.** residues). A plot of the ARI across different date cutoffs in Fig. 7 shows that our predicted consensus pockets are very stable for nsp5 over different input data but more variable for nsp12 and Spike. Images on the right side of Fig. 7 illustrate the overlap between pairs of clusterings and thus provide a physical context for interpreting the ARI scores in the plot. Although the ARI is noticeably lower for the earliest two date cutoffs for nsp12, it covers a more modest range over the rest of the dates and is relatively consistent from 06/23/2020 through 10/05/2021. While the ARI for Spike drops consistently from more recent to older date cutoffs, our process still identifies the same literature-confirmed pocket at every date cutoff except 07/09/2019 and 03/10/2020, when data on SARS-CoV-2 was more limited. Since more data on a new pathogen will be available at later dates, we would naturally expect that predictions from our data-driven approach will improve over time and that there will be some moderate time-sensitivity in our results. Yet even early on in the pandemic, when data on SARS-CoV-2 was limited, there seems to be enough data from other related organisms for our framework to detect comparable binding pockets. Together, the findings from these internal validation tests lend support for the effectiveness of both our structural modeling methods and consensus pocket identification pipeline.

**Figure 7:**
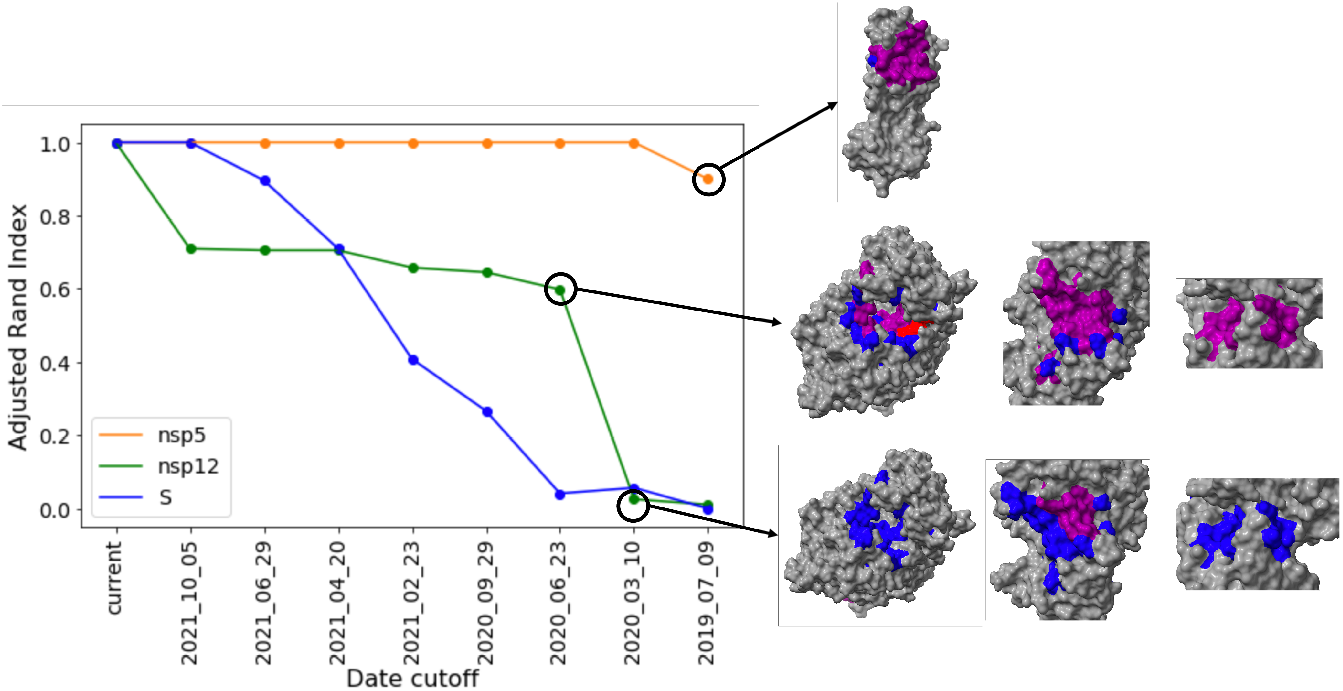
On the left, a plot of the adjusted rand index (ARI) for the consensus pockets in nsp5, nsp12, and Spike as a function of date cutoff in the PDBspheres data. The GDC cutoff is held fixed at 60. An ARI of one indicates that two sets of consensus pockets are identical, while an ARI of zero indicates that the consensus pockets in each set have no residues in common. The reference consensus pockets are those associated with a GDC cutoff of 60 and a current date cutoff. On the right, images of the overlap between the reference clustering and three comparison clusterings noted with a black circle. Residues colored blue, red, and purple appear only in the consensus pockets of the reference point, only in the consensus pockets of the point of comparison, or in both, respectively. The top, middle, and bottom rows illustrate high, medium, and low ARI values.

To further assess the robustness of our methods, we examine the relationship between the GDC metric in the PDBspheres data and the source organisms of the underlying PDB experimental complexes. For each organism category, we record the GDC scores of all protein-ligand binding conformations in the PDBspheres data with a source organism belonging to that category and plot the distribution of scores. In the case of PDB complexes with more than one source organism, we count the GDC score towards all source organisms, since our approach does not distinguish which parts of a PDB complex originate from which source organism. As shown in Fig. 8, for source organisms SARS-CoV-1 and SARS-CoV-2, the distribution of GDC scores has one peak between 90 and 95 with a long, thin tail at lower GDC values, indicating a high level of structural similarity with the SARS-CoV-2 proteins of interest, as expected. The distribution for other viruses similarly peaks around 90-95, but it carries more density across lower GDC values than do the distributions for SARS-CoV-1 and −2. In contrast, for other organism classifications, with the exception of insects, the distribution either peaks around 60-70 or is more uniform across the range of GDC values, signaling a lower degree of structural similarity with SARS-CoV-2 proteins. The peak at high GDC in the insect distribution is an artifact and can be attributed to a single PDB complex with components from both SARS-CoV-2 and the saltans group. As noted above, the GDC score for this complex counts towards both the SARS-CoV-2 category and the insect category, even though the high score is almost certainly a result of the SARS-CoV-2 component and its high degree of similarity with the SARS-CoV-2 nsp3 structural model. These results confirm that, as expected, the less closely related an organism is to SARS-CoV-2, the less structurally similar its pockets are to those in SARS-CoV-2.

**Figure 8:**
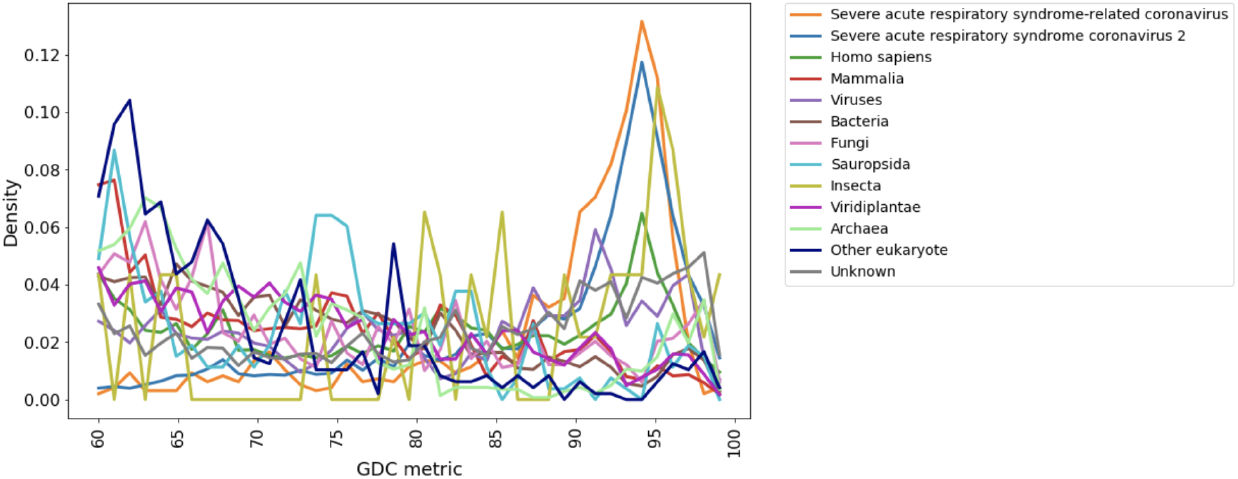
Distributions of the GDC metric for PDBspheres structural models from various organism classifications.

The trend between the amount of PDBspheres data and time is also informative. As expected, the number of protein-ligand pairs in PDBspheres with a GDC score above 60 increases over time for all SARS-CoV-2 proteins (Fig. S5, left panel). We note that the number of proteinligand pairs grows faster and reaches a higher point for Spike than for other proteins simply because there are more unique protein templates for Spike, as seen in the right panel of Fig. S5. We can conclude that pockets detected through PDBspheres are stable over time. That is, once PDBspheres detects a pocket, it continues to detect the pocket even as additional, newer data becomes available. Therefore, we can be confident in the quality of the PDBspheres structural models even early in a pandemic when experimental structural data on a new pathogen is limited.

Our method is unique not only in its emphasis on a comprehensive data-driven search of protein structure conservation across the entire PDB, but also in its pocket characterization. Indepth analysis of the composition of the consensus pockets in terms of their underlying PDBspheres models provides additional context for our results, highlighting one aspect of its characterization capability. For each consensus pocket, we determine which PDBspheres models correspond to the protein-ligand conformations that involve that pocket. For each relevant proteinligand conformation, we include as contributing models both the model for the target protein of interest and the model for the protein with the observed or predicted ligand binding. For each contributing model, we then retrieve from PDB the source organism for the experimental measurement from which the model was generated. If a single model corresponds to more than one organism, all organisms are counted for that model. Pie charts of the contributions from various organism classifications allow for straight-forward visual comparisons across different data filters, as shown in Fig. 9. Observing the evolution of organism composition over time, we see that, by definition, there is no contribution from SARS-CoV-2 in consensus pockets identified from pre-pandemic data and that the SARS-CoV-2 contribution grows over the course of the pandemic as more data on the new virus is collected. The source organism composition of consensus pockets could be utilized as a preliminary means of narrowing in on pockets that are more highly conserved across SARS-CoV-2 and other viruses — pockets that may hold more promise as targets for broad-spectrum antivirals.

**Figure 9:**
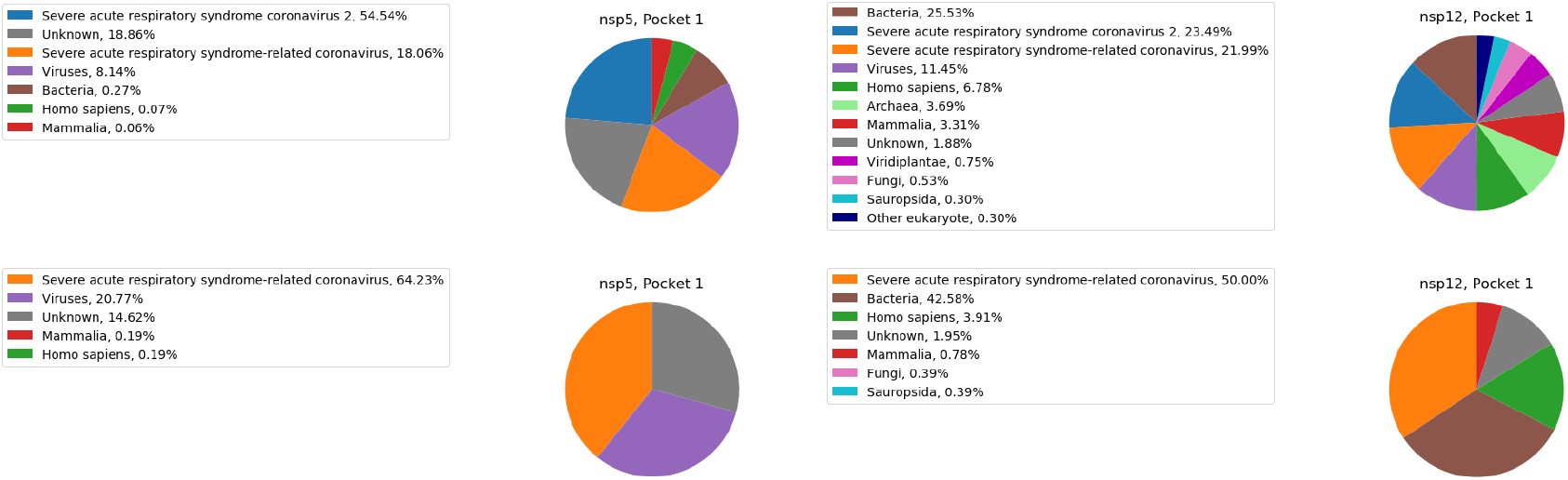
Composition of consensus pockets in terms of the source organisms for the underlying PDBspheres models, shown on a log scale. The left and right columns show corresponding consensus pockets for nsp5 and nsp12, respectively, using currently available data (top row) and pre-pandemic data (bottom row). Pockets were numbered consistently by inspection between iterations of our pipeline across different date cutoffs.

Based on this reasoning, we examine the organism composition profiles of the consensus pockets from a range of SARS-CoV-2 proteins and calculate a simple ranking of the pockets’ degree of structural conservation. The score *S* for each pocket *p* is determined by the size of the viral contribution (including SARS-CoV-1, SARS-CoV-2, and other Viruses) relative to the overall contribution from known organisms and is given by

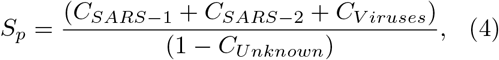

where *C_i_* is the fraction of pocket p’s pie chart that component *i* makes up. Pockets are ordered from highest to lowest score, with higher scores indicating a greater specificity for viral sites in comparison to sites from humans or other organisms. Fig. 10 shows the highest scoring pocket for each protein and the range of composition profiles observed across consensus pockets. Proteins without enough PDBspheres data for determination of consensus pockets are excluded from this analysis. Preferred profiles, like those for nsp5-pocket1 and nsp12-pocket2, tend to have fewer different contributing organisms, a larger share of the composition derived from viruses, and minimal contribution from humans (which could, though not necessarily, indicate a potential for undesirable off-target interactions). We note that it is possible for a pocket to have a larger contribution from SARS-CoV-1 than SARS-CoV-2, as is the case with nsp14-pocket1, if the contributing models derived from SARS-CoV-1 are predicted to bind with a larger number of ligands and therefore appear more frequently in the PDBspheres data. Comparing the composition of all consensus pockets identified for Spike, nsp5, and nsp12, we find that overall, with some exceptions, these three preferable features are more common in the profiles for nsp5 and nsp12 than in Spike, as shown in Fig. S6. Together, these observations suggest that the consensus pockets in nsp5 and nsp12 are more structurally similar to binding pockets in other viruses, which may help explain why our method seems to perform better for nsp5 and nsp12 than for Spike. Indeed, RNA Dependent RNA Polymerases like nsp12 are known to be highly conserved across different viruses. ^37^ Given that half of our nsp12 pockets map to the Pfam RdRp_1 family, the structural similarity we detect between the pockets in nsp12 and other viruses is expected.

**Figure 10:**
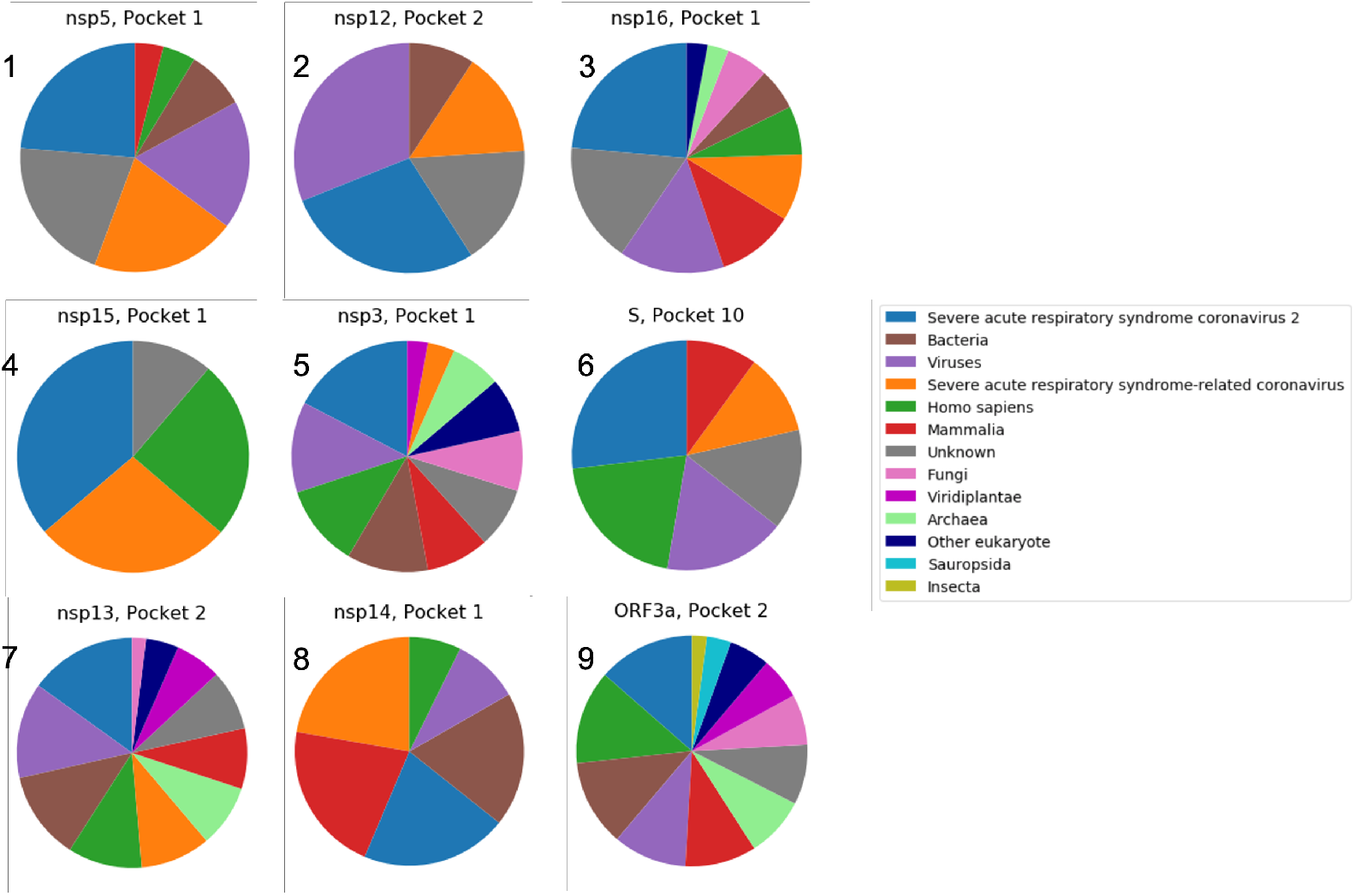
Composition of consensus pockets in terms of the source organisms for the underlying PDBspheres models, shown on a log scale. The top pocket for each protein is shown, as determined by the size of the viral contribution (SARS-CoV-1, SARS-CoV-2, plus other Viruses) relative to the overall contribution from known organisms. Composition profiles are numbered from most to least preferred.

Exploring the PDBspheres data from a ligand-centric perspective provides further insight into patterns of protein-ligand binding. Clustering of the ligands that bind to each consensus pocket reveals groups of ligands with clearly identifiable and distinct features, such as functional groups, molecular size, and geometry. Together these features define a unique chemotype for each cluster (as seen in the example ligands for nsp5-pocket1 on the right side of Fig. 11), and the collection of chemotypes for a consensus pocket can be used to characterize it. The following aspects of the ligand clustering silhouette plots (Fig. S2), which are typical of our results, indicate separation between ligand clusters and support their validity: mostly positive individual silhouette coefficients for members of each cluster, a positive average silhouette coefficient, and an average coefficient that crosses each of the clusters.

**Figure 11:**
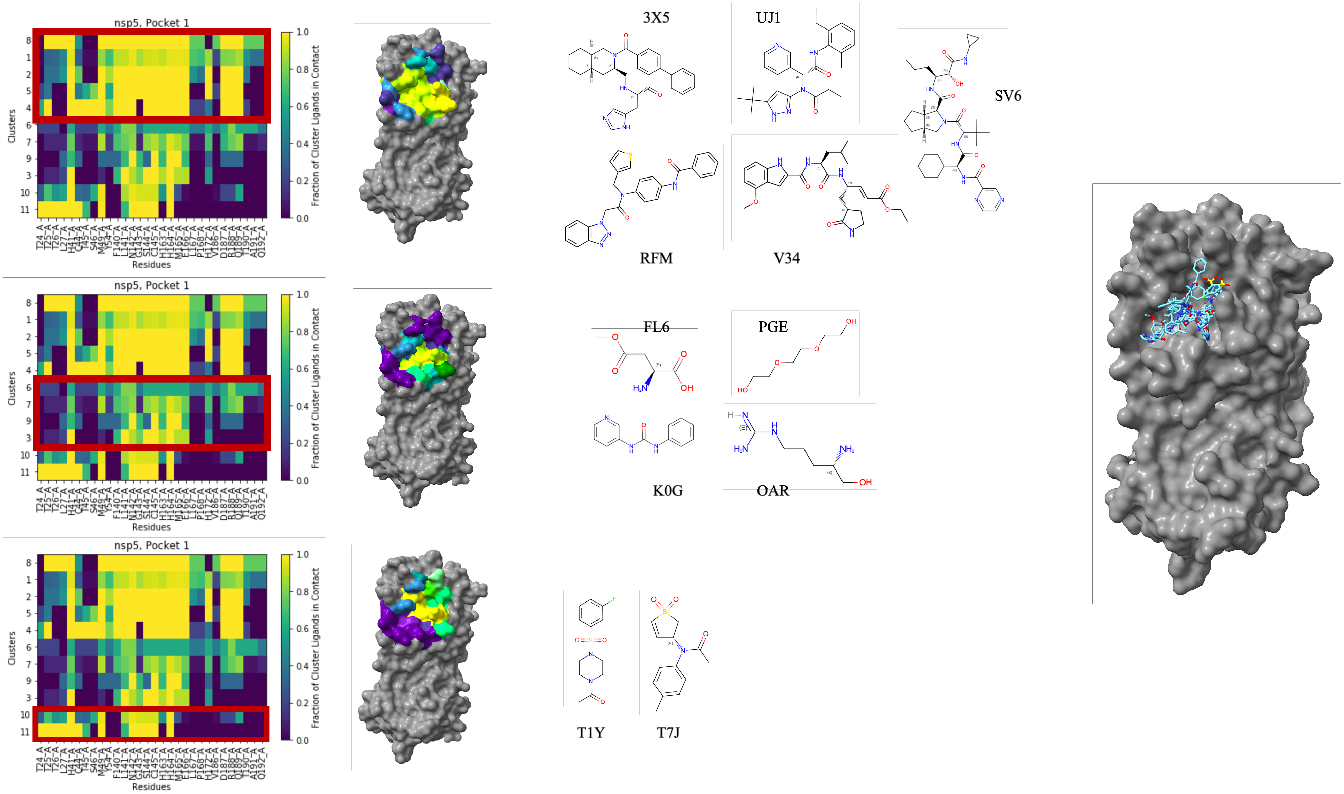
On the far left, heatmaps of the fraction of ligands in each cluster with which residues in nsp5-pocket1 come into contact. In the center left, 3D images of the surface of nsp5 with pocket residues colored by the average fraction within the supercluster designated in the red box in the corresponding heatmap. In the center right, images of one representative molecule from each ligand cluster in the corresponding supercluster. Molecules are labeled by their three-character PDB ligand ID. SMILES strings for each molecule are provided in Supporting Information. On the far right, an image of nsp5-pocket1 with all eleven representative molecules superimposed.

Analysis of the fraction of ligands in each cluster that contact each residue in a given pocket enables us to locate subregions of the pocket that correspond to specific chemotypes. As seen in the heatmaps on the left side of Fig. 11, we find that different ligand clusters have similar patterns of contact fractions for the residues in the pocket. We group the ligand clusters according to the pattern of contact fractions and calculate the average contact fraction for each residue over the clusters in the super-cluster. This average contact fraction determines the color of the residue in the 3D images of the protein surface, shown in the center of Fig. 11. As seen on the right side of Fig. 11, we observe that larger subregions correspond to larger chemotypes, a physically reasonable result. Our ligand clustering method thus provides a finer resolution view of our consensus pockets and the ability to hone in on smaller regions of interest for specific chemotypes.

Results compiled from several experimental assays^1^ that tested activity against nsp5 lend support for our approach to ligand clustering. Positive hits were based on multi-concentration IC50 measurements, and negative hits were subjected to single concentration screens at values ranging from 10 —100*μM* depending on the screen. Additional details on the assays are described in Ref. 38 and 39. Using Tanimoto similarity, we compare the structure of the experimental compounds to the structure of the ligands in each ligand cluster for pocket 1. For each experimental compound, we determine the ligand cluster to which it is closest on average. Then for each cluster, we compare the distributions of average Tanimoto distances for the positive and negative compounds assigned to that cluster. Fig. 12 displays these distributions as well as the results of a t-test on the mean values, omitting clusters for which no experimental compounds were assigned or too few compounds were assigned to statistically compare the positive and negative distributions. P-values above or below the significance level of *α* = 0.05 indicate that the mean Tanimoto distances for positive and negative compounds are statistically equivalent or different, respectively. While the positive and negative compounds are roughly the same distance from two of the clusters, for four other clusters, the positive compounds are on average closer and thus more similar in structure than are the negative compounds. The finding that experimentally observed positive compounds align more closely with our clusters than do negative compounds affirms that our ligand clustering method detects meaningful chemical features associated with effective binding. Though not all ligand binding leads to biological impact, comparing such histograms across ligand clusters and identifying those with the greatest degree of separation between the positive and negative distributions could signal which chemotypes deserve further attention and study. Following identification of highly conserved consensus pockets, assessment of the drug likeness of site-specific ligands, either through standard drug-likeness tests or by calculating average Tanimoto similarity against a library of known drugs, may serve as an additional means for identifying promising broad-spectrum drug targets. More detailed simulation and/or experiments could then be conducted on the chemotypes of interest to investigate whether their binding causes functional effects.

**Figure 12:**
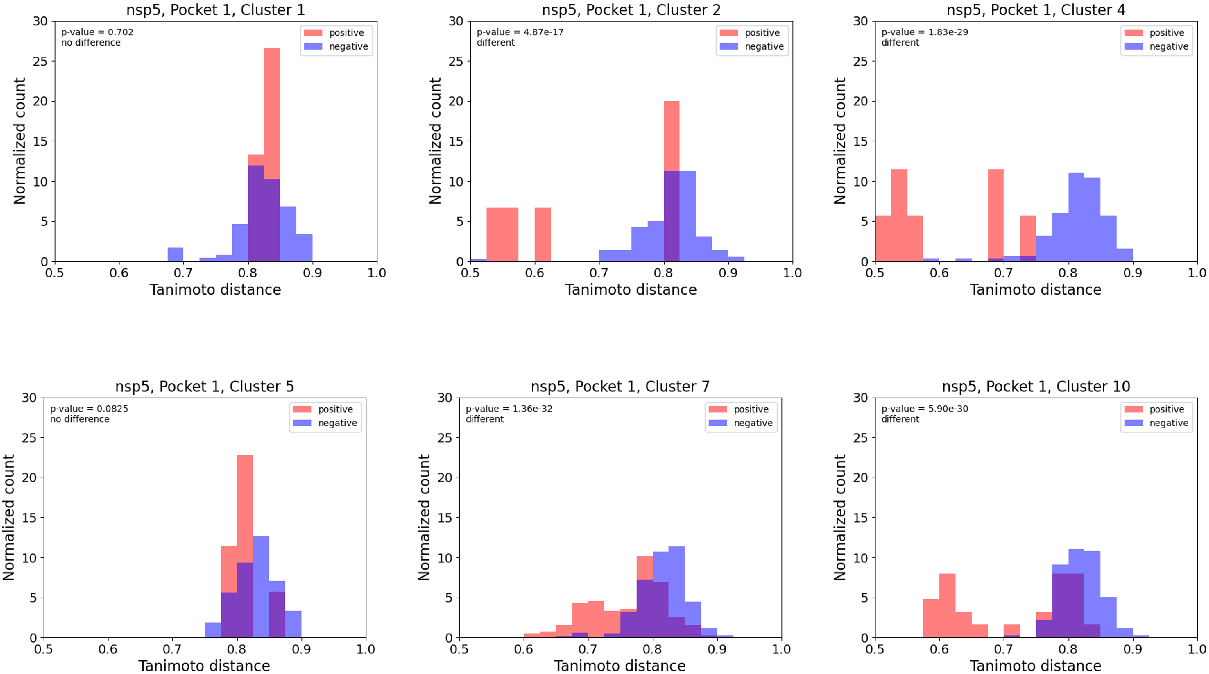
Normalized histograms of the average Tanimoto distance between experimentally-tested compounds and their closest ligand cluster. The distributions for positive and negative compounds are shown in red and blue, respectively. P-values shown on each plot are for the associated t-test comparing the means of the positive and negative distributions.

While the ligands in some clusters are insignificant (for example, molecules used in buffer solutions in experimental structural measurements in PDB), those in other clusters include molecules that are relevant for drug development. Having omitted PDB data for known SARS-CoV-2 drugs when running our pipeline, we compare the chemical structure of one such drug to our ligand clusters. PF-07321332, or Nirmatrelvir, is a component of the antiviral Paxlovid and is associated with nsp5 in the PDB templates 7SI9 and 7VH8 from Fig. 6. As with the experimental compounds in Fig. 12, we calculate the Tanimoto distance between Nirmatrelvir and each ligand in each ligand cluster for nsp5. We find that, based on the mean Tanimoto distance from each cluster, Nirmatrelvir is closest to cluster8, which contains two known antivirals for Hepatitis C: Boceprevir and Telaprevir.

The fact that our process links known drugs for two different viruses suggests that our approach offers an avenue for identifying chemical features or substructures that hold potential for broad-spectrum drug development. In light of this observation, we put forward the following strategy for identifying promising chemical substructures from our results. First, look for known drugs among the ligand clusters. Second, look at the unclustered ligands that are near the clusters containing known drugs. Unclustered ligands are ones deemed by the DBSCAN algorithm to be “outliers” or “noisy” points due to their relative isolation from other ligands. Yet unclustered does not necessarily imply irrelevant; some drugs or drug-like molecules may have unique features that differentiate them from the clustered ligands. Finally, check chemical databases for compounds that are not found in PDB but are similar to the drug-containing clusters and/or the nearby unclustered ligands. Within the set of molecules of interest, (clusters containing known drugs, nearby unclustered ligands, and related non-PDB molecules), common chemical features or substructures may serve as starting points for further refinement and optimization in the drug discovery process.

## 5 Conclusion

Here we present a new approach for detecting and characterizing relevant binding pockets in the proteins of novel pathogens and for uncovering the site-specific, core chemical substructures associated with protein-ligand binding. Our novel computational method is unique in its straightforward, data-driven use of purely experimental structural data, its strength in identifying structurally conserved binding pockets, and its comprehensive analysis and characterization of both pockets and associated ligands. We combine molecular level detail from protein structural modeling with machine learning clustering techniques and cheminformatics tools in a predictive pipeline that is fast, streamlined, and broadly applicable. While the quality of available protein structural models is a limiting factor, we account for this potential issue by considering a quantitative measure of model quality, the GDC score. Only structural models above a desired quality are included in our downstream analysis. Our framework can be deployed as soon as a new pathogen is discovered and can generate results in a matter of days. We find that it performs as intended even when experimental structural data on a new pathogen is limited, and our predictions should improve over time as more data is collected. With a basis in experimental data and few parameters, our process is straightforward and can be readily extended to new pathogens, provided their genomic sequence is available or sufficiently accurate homology models can be created. Indeed, its applicability holds more generally for any class of proteins with structurally conserved pockets, such as kinases and G-protein-coupled receptors.

In this paper, we focus on SARS-CoV-2 as a case study for explaining our system and assessing its performance. Our predictions of consensus binding pockets in the nsp5, nsp12, nsp14, nsp15, nsp16, and Spike proteins compare favorably with those already published in the literature, including some detected through experiment and others via computational methods. While our consensus pocket identification method is somewhat sensitive to the input data, modest variations like those we observe are to be expected. Even though the similarity metric declines going back in time, in many cases we still identify the pockets reported in other published papers.

As expected, we observe that the more closely related an organism is to SARS-CoV-2, the more structurally similar its binding pockets are to those in SARS-CoV-2. Analysis of the composition of source organisms of our consensus binding pockets reveals that nsp5 and nsp12 are more structurally conserved viral proteins than is Spike.

Through our ligand clustering method, we identify characteristic chemotypes for each consensus pocket as well as the subregions of the pocket with which each chemotype interacts. Significantly, we find that positive experimental hits against nsp5 are on average more similar in structure to our ligand clusters than are the negative compounds from the screening – evidence that the core chemical components we detect are in fact relevant.

Our results align well with current thinking on key goals in the search for broad-spectrum antiviral drug targets and candidates. A SARS-CoV-2 binding pocket may hold more promise as a broad-spectrum antiviral drug target 1) the more similar its structure is to that of other viral binding pockets and 2) the more drug-like the molecules that bind. Our pipeline offers a means of assessing these two factors, through source organism composition analysis and ligand clustering. While nsp5 and nsp12 have received more attention to date, our results affirm other findings of conserved pockets in nsp3, nsp13, and nsp16^17,18^ and suggest that these proteins should be studied further, as additional data would provide more clarity and confidence. Overall, our system lays the groundwork for a rapid response tool that can quickly identify promising drug targets and target-specific drug compound components when faced with an emerging pathogen, and ongoing work will enhance these capabilities.

## 6 Data and Software Availability

The complete PDBspheres library of data is available at https://proteinmodel.org/AS2TS/PDBspheres. Code and input data for the residue and ligand clustering procedures is located at https://github.com/llnl/TargetID. All other software is open source and cited throughout the paper.

## 7 Supporting Information

Additional data and figures in support of our methods, a list of the constituent residues of the consensus pockets identified for each SARS-CoV-2 protein, and SMILES strings for the molecules in Fig. 11

## 8 Acknowledgements

Molecular graphics created with UCSF ChimeraX, developed by the Resource for Biocomputing, Visualization, and Informatics at the University of California, San Francisco, with support from National Institutes of Health R01-GM129325 and the Office of Cyber Infrastructure and Computational Biology, National Institute of Allergy and Infectious Diseases. See Ref. 40 and Ref. 41 for more details. Venn diagrams created with InteractiVenn (http://www.interactivenn.net/). See Ref. 42 for more details. Financial support for this work is provided by the Laboratory Directed Research and Development program at Lawrence Livermore National Laboratory (20-ERD-062).

