## Supporting Information for "A Computational Pipeline to Identify and Characterize Binding Sites and Interacting Chemotypes in SARS-CoV-2"

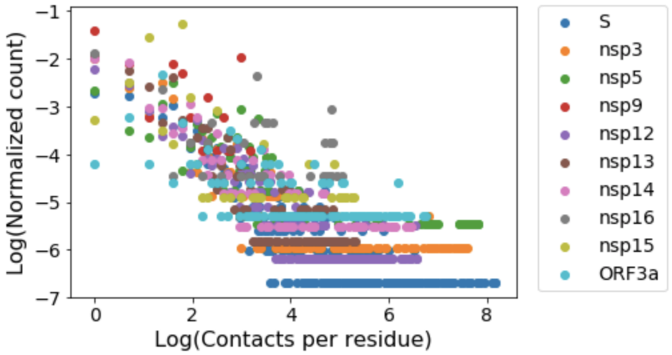


Figure S1: Distribution of contacts per residue in the PDB Spheres data for each SARS-CoV-2 protein.

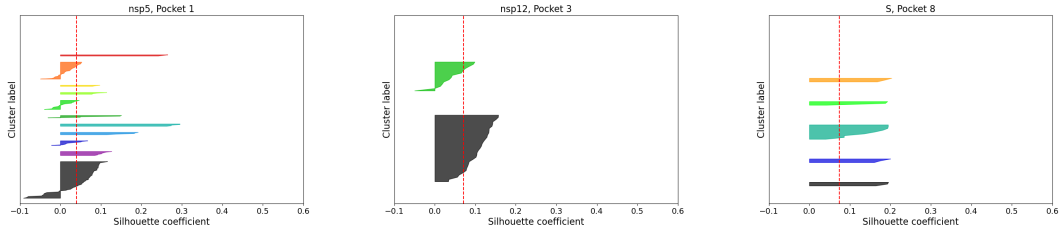


Figure S2: Sample silhouette plots for the ligand clusters of nsp5-pocket1 (left), nsp12-pocket3 (middle), and Spike-pocket8 (right) indicate reasonable separation between ligand clusters based on the following three features: mostly positive individual silhouette coefficients for members of each cluster, a positive average silhouette coefficient, and an average coefficient that crosses most of the clusters.

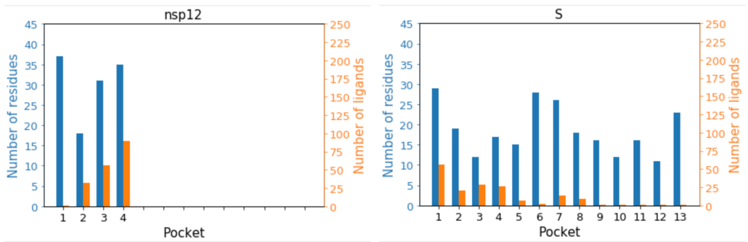


Figure S3: Bar charts show the number of constituent residues (blue) and number of ligands (orange) that bind at least 50% of the residues for the consensus pockets in nsp12 (left) and Spike (right).

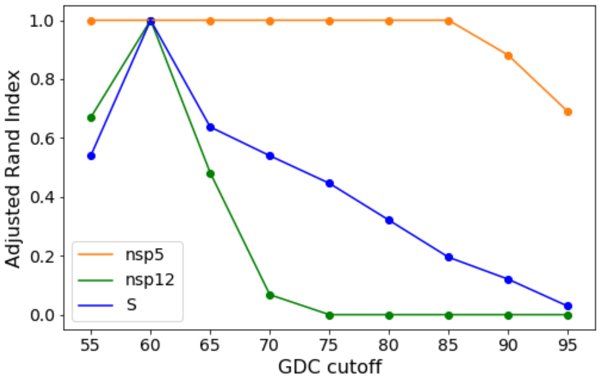


Figure S4: Plot of the adjusted rand index (ARI) for the consensus binding sites in nsp5, nsp12, and Spike as a function of GDC cutoff in the PDB Spheres data. An ARI of one indicates that two sets of binding sites are identical, while an ARI of zero indicates that the binding sites in each set have no residues in common. The reference binding sites are those associated with a GDC cutoff of 60 and a current date cutoff. The date cutoff is held fixed at "current" in this plot.

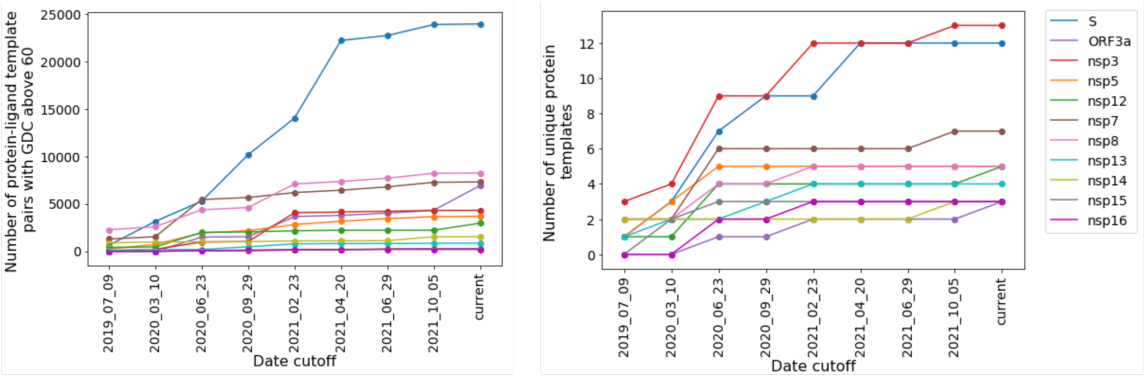


Figure S5: Number of protein-ligand pairs with a GDC above 60 (left) and unique protein templates (right) in the PDB Spheres data for each SARS-CoV-2 protein as a function of time.

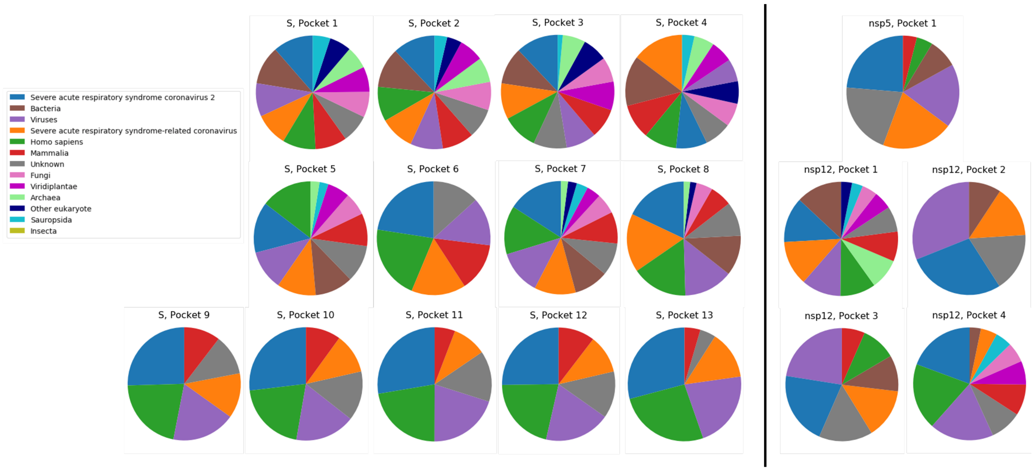


Figure S6: Composition of consensus pockets in terms of the source organisms for the underlying PDBspheres models, shown on a log scale, for all pockets identified in Spike (left side), nsp5 (right side, top) and nsp12 (right side, center and bottom).

| Pocket Number | Residues |
|---|---|
| 1 | A225, D226, I227, V228, A242, A243, N244, K248, H249, G250, G251, G252, V253, A254, A256, P329, L330, L331, S332, A333, G334, I335, F336, G337, P340, A358, V359, F360, D361, L364 |
| 2 | L214, L216, I222, D339, I341, H342, R345, K362, N363, Y365, D366, K367, V369, S370, S371, F372, L373, E374, K376, S377, E378 |
| 3 | W851, N854, N855, C856, Y857, K902, E906, L907, G908, D909, R911, E912, M953, G954, A991, P992, P993, Y1009, T1010, G1011, N1012, Y1013, Q1014, C1015, G1016, H1017, Y1018, K1019, D1031, T1046 |

Table S1: Consensus binding sites for nsp3. All residues belong to the A chain.

| Pocket Number | Residues |
|---|---|
| 1 | C93, P94, E180, R183, Q184, L186, L187, K188, T189, V190, Q191, F192, C193, D194, A195, R197, N198, G214, W216, V231, Y237, Y238, H256, V257, D258, T259, D260, L282, F283, D284, R285, Y286, F287, K288, Y289, W290, D291 |
| 2 | N496, K500, S501, K511, N543, L544, Y546, V557, A558, G559, R569, K577, F594, Y595, G683, D684, A685, Y689 |
| 3 | D499, R513, K545, I548, R555, A580, Y619, C622, D623, T680, S682, T686, T687, A688, N691, L758, S759, D760, D761, C813, S814, Q815, R836, A840, D845, K849, E857, R858, S861, L862, D865 |
| 4 | V30, V31, Y32, R33, A34, F35, K50, N52, C53, R55, Y69, V71, K73, H75, N79, E83, A97, K98, H99, R116, Q117, L119, T120, K121, Y122, T123, D126, D208, N209, Q210, D211, Y217, D218, F219, G220 |

Table S2: Consensus binding sites for nsp12. All residues belong to the A chain.

| Pocket Number | Residues |
|---|---|
| 1 | V6, A18, C19, I20, R21, R22, P23, F24, H39, A110, R129, L132, F133, E136, T137, A140, T141, T144, V232, M233, P234, L235, S236 |
| 2 | W114, D119, Y120, I121, L122, A123, N124, T125, K131, A134, A135, L138, K139, E142, A379, T380, N381, Y382, D383, A407, P408, R409, T410, Y421, F422, N423 |

Table S3: Consensus binding sites for nsp13. All residues belong to the A chain.

| Pocket Number | Residues |
|---|---|
| 1 | D90, V91, E92, G93, H95, P141, F146, L149, H268, D273 |
| 2 | D301, E302, L303, K304, I305, N306, A307, A308, R310, K311, H314, K318, C340, V341, P342, Q343, A344, W348, N422, N489, L493, Y494, D496, A497, M500 |
| 3 | Y465, V466, L468, K469, S470, Y491, R492, L495, N499, S503 |

Table S4: Consensus binding sites for nsp14. All residues belong to the A chain.

| Pocket Number | Residues |
| --- | --- |
| 1 | H234, Q244, L245, G246, G247, H249, K289, V291, C292, S293, W332, E339, T340, Y342, P343, K344, L345 |

Table S5: Consensus binding sites for nsp15. All residues belong to the A chain.

| Pocket Number | Residues |
| --- | --- |
| 1 | N43, K46, Y47, H69, G71, A72, G73, S74, A79, P80, G81, D99, L100, N101, G113, D114, C115, D130, M131, Y132, D133, P134, F149, K170, N198 |

Table S6: Consensus binding sites for nsp16. All residues belong to the A chain.

| Pocket Number | Residues |
|---|---|
| 1 | Y756, F759, Q762, L763, A766, I770, T961, Q965, S968, N969, F970, G971, V991, D994, R995, T998, G999, L1001, Q1002, S1003, Q1005, T1006, Y1007, V1008, T1009, Q1010, L1012, I1013, E1017 |
| 2 | N764, R765, L767, T768, G769, V772, E773, K776, N777, E780, A1015, A1016, R1019, A1020, N1023, L1024, T1027, R1039, F1042 |
| 3 | E725, I726, L727, P728, K947, D950, V951, Q954, Q957, R1014, I1018, S1021 |
| 4 | I973, S974, S975, V976, L977, N978, D979, I980, L981, S982, L984, D985, V987, E988, Q992, D1163, D1165 |
| 5 | V826, T827, L828, A829, D830, A831, L849, L948, Q949, V952, N953, A956, P1057, H1058, G1059 |
| 6 | C336, P337, F338, G339, E340, V341, F342, N343, I358, A363, Y365, V367, L368, Y369, N370, S371, A372, S373, F374, F377, L387, F392, V395, I434, W436, L513, F515, V524 |
| 7 | T716, N717, F718, T719, I720, S721, V722, T723, T724, N919, K921, L922, N925, Q926, N928, S929, A930, G932, K933, I934, D936, S937, S940, A1070, Q1071, F1109 |
| 8 | I794, K795, D796, G799, F800, N801, F802, S803, Q804, I805, F817, I818, F927, I931, Q935, F1052, L1063, V1065 |
| 9 | V11, V120, N122, A123, T124, N125, V127, K129, E154, F157, Y160, E169, V171, Q173, P174, L176 |
| 10 | T108, T109, T114, Q115, N196, G199, G232, I233, N234, I235, T236, R237 |
| 11 | A27, Y28, T29, N30, F32, F59, S60, N61, V62, T63, D215, Q218, V267, Y269, P631, R634 |
| 12 | K1073, F1075, N1098, G1099, T1100, H1101, W1102, F1103, Y1110, P1112, I1114 |
| 13 | R346, F347, A348, W353, N354, R355, P426, F429, T430, R454, R457, K458, S459, N460, L461, K462, P463, F464, E465, R466, D467, E471, S514 |

Table S7: Consensus binding sites for Spike. All residues belong to the A chain.

| Pocket Number | Residues |
|---|---|
| 1 | $I37_A$, $A39_A$, $S40_A$, $L41_A$, $P42_A$, $W45_A$, $V48_A$, $G49_A$, $L52_A$, $V55_A$, $F56_A$, $L73_A$, $G76_A$, $V77_A$, $V80_A$, $L83_A$, $L84_A$, $L86_A$, $F87_A$, $V90_A$, $Y91_A$, $L94_A$, $F120_A$, $I124_A$, $R126_A$, $L127_A$, $L139_A$, $L140_A$ |
| 2 | $L46_A$, $I47_A$, $V50_A$, $A51_A$, $L53_A$, $A54_A$, $Q57_A$, $S58_A$, $S60_A$, $K61_A$, $I63_A$, $L65_A$, $S74_A$, $K75_A$, $H78_A$, $F79_A$, $C81_A$, $N82_A$, $L85_A$, $V88_A$, $T89_A$, $S92_A$, $H93_A$, $L96_A$, $F105_A$, $L106_A$, $L108_A$, $Y109_A$, $A110_A$, $L111_A$, $V112_A$, $Y113_A$, $F114_A$, $L115_A$, $Q116_A$, $S117_A$, $N119_A$, $R122_A$, $I123_A$, $D142_A$ |
| 3 | $S40_B$, $L41_B$, $P42_B$, $W45_B$, $L46_B$, $I47_B$, $V48_B$, $G49_B$, $V50_B$, $L52_B$, $L53_B$, $F56_B$, $W69_B$, $L73_B$, $K75_B$, $G76_B$, $V77_B$, $H78_B$, $F79_B$, $V80_B$, $C81_B$, $N82_B$, $L83_B$, $L84_B$, $L85_B$, $L86_B$, $F87_B$, $V88_B$, $T89B$, $V90_B$, $Y91_B$, $H93_B$, $L94_B$, $Y109_B$, $V112_B$, $Y113_B$, $Q116_B$, $L127_B$, $L139_B$, $X403_C$ |

Table S8: Consensus binding sites for ORF3a. The subscript indicates the chain on which the residue is located.
